# An integrated DNA interactome and transcriptome profiling reveals a PU.1/enhancer RNA-mediated Feed-forward Regulatory Loop Regulating monocyte/macrophage development and innate immune functions

**DOI:** 10.1101/2025.02.19.638695

**Authors:** A S M Waliullah, Kevin Qiu, Barbara Dziegielewska, Minh L. Tran, Nicholas N. Nguyen, Leran Wang, Andrew Pan, Natalie Segovia, Simone Umarino, Junyan Zhang, Tuan M. Nguyen, Jeffrey Craig, Daniel G. Tenen, Bon Q. Trinh

## Abstract

High expression of the myeloid master ETS transcription factor PU.1 drives the development of monocyte/macrophage (Mono/MΦ), a crucial cellular component of the innate immune system. Disruptions in normal expression patterns of PU.1 are linked to a variety myeloid malignancy and immune diseases. It is evidenced that PU.1 binds to and modulates enhancers of several myeloid genes. While noncoding RNAs transcribed from noncoding genes at the enhancers are increasingly reported to be involved in enhancer regulation, the crosstalk between PU.1 and noncoding RNAs in enhancer-mediated myeloid gene regulation in Mono/MΦ differentiation and immune response has not been systematically investigated. In this study, we interrogated the PU.1-mediated transcriptome and cistrome with our comprehensive collection of putative and verified enhancers. Among a repertoire of noncoding genes present at PU.1-bound enhancers, we discovered that PU.1 acts as a potent transcription factor inducer of the noncoding RNA *LOUP*, which we previously identified as an RNA inducer of PU.1. The genomic region within the *LOUP* locus occupied by PU.1 is characterized by the epigenetic features of a myeloid-specific super-enhancer. Targeted disruption of the PU.1-binding motifs resulted in the downregulation of *LOUP* promoter activity. Depletion of *LOUP* reduced the expression of Mono/MΦ cell markers as well as the transcriptional program associated with Mono/MΦ differentiation Mono/MΦ innate defense mechanisms, including phagocytosis, antimicrobial activity, and chemoattractant cytokine production. *LOUP* induces Mono/MΦ phagocytic activities. Collectively, our findings indicate that PU.1 and enhancer RNA *LOUP* are biomolecular components of an unidentified feed-forward loop that promotes their mutual expression, contributing to Mono/MΦ differentiation and innate immune functions. The identification of the PU.1/*LOUP* regulatory circuit provides valuable insights into the mechanisms underlying cell-type and gene-specific enhancer activity and Mono/MΦ biology, as well as significant implications for advancing our understanding of immune diseases and myeloid malignancies.

## INTRODUCTION

Monocytes/macrophages (Mono/MΦ) are essential cellular constituents of the myeloid cell lineage that play a crucial role in the body’s defense against infections (De Kleer, Willems et al. 2014, Ginhoux and Jung 2014). Mono/MΦ development is ensured by predominant levels of the myeloid master ETS transcription factor (TF) PU.1 (also known as SPI1 or SFPI1) (Chen, Zhang et al. 1995, Korczmar, Bookstaver et al. 2024). PU.1 is essential for the development of multipotential myeloid precursor cells and is required for terminal Mono/MΦ differentiation (Tenen 2003). Reduced expression of PU.1 disrupts myeloid cell differentiation, resulting in the development of acute myeloid leukemia (AML) (Cook, McCaw et al. 2004, Rosenbauer, Wagner et al. 2004). Accumulating evidence also demonstrates the role of PU.1 in Mono/MΦ immune functions. For instance, PU.1 promotes mature macrophage and governs innate immune responses (Karpurapu, Wang et al. 2011, Qian, Deng et al. 2015, Fischer, Walter et al. 2019). Additionally, PU.1 is involved in pathological conditions involving Mono/MΦ, such as inflammation related to asthma and allergies (Qian, Deng et al. 2015, Fischer, Walter et al. 2019). While the role of PU.1 in Mono/MΦ development is well-studied, its role and mechanisms in Mono/MΦ functions remain poorly understood. Nevertheless, to infer its critical role as a TF, it is imperative to understand how PU.1 regulates gene expression in Mono/MΦ development and innate immune functions.

A number of studies indicate that PU.1 binds to and modulates enhancers of myeloid genes. For instance, PU.1 binds a +37 kb enhancer that induces the expression of the myeloid gene *Cebpa* in murine cells (Cooper, Guo et al. 2015). PU.1 also binds and stimulates the myeloid-specific −2.7 kb enhancer to drive the expression of chicken lysozyme (Faust, Bonifer et al. 1999). Furthermore, PU.1 binds the conserved −17 kb enhancer in humans (−14 kb in mice), known as the upstream regulatory element (URE) (Li, Okuno et al. 2001, Ebralidze, Guibal et al. 2008, Staber, Zhang et al. 2013). Abrogation of the PU.1 binding site within the URE impairs the formation of the URE-promoter chromatin loop, leading to downregulation of *PU.1* expression in myeloid cells (Chen, Ray-Gallet et al. 1995, Leddin, Perrod et al. 2011, Staber, Zhang et al. 2013). The URE is part of an ∼8 kilobase *<u>P</u>U.1* <u>ci</u>s-regulatory <u>e</u>lement <u>c</u>luster (PCREC) that displays molecular features of myeloid-specific enhancers (Qiu, Vu et al. 2024). Moreover, introducing PU.1 into non-hematopoietic cells where it is not normally expressed provides PU.1 access to chromatin and enables chromatin remodeling, suggesting that PU.1 acts as a pioneering factor that establishes an active enhancer state (Barozzi, Simonatto et al. 2014, Ha, Cho et al. 2019). However, a comprehensive survey of PU.1-bound enhancers (PU.1-Es) in myeloid cells—particularly their role in Mono/MΦ development and function—has not yet been performed.

Increasing evidence has indicated the important role of noncoding genes (NGs) in enhancer activities. NGs constitute a significant portion of the mammalian genome, including genes that lack protein-coding sequences and a subset of pseudogenes that resemble functional genes. These genes are transcribed but not translated, giving rise to numerous noncoding RNAs (ncRNAs) (Costa 2008, Mercer, Dinger et al. 2009, Ponting, Oliver et al. 2009, Groen, Capraro et al. 2014). To date, only a few ncRNAs have been precisely mapped and functionally characterized (Uszczynska-Ratajczak, Lagarde et al. 2018), leaving the majority poorly annotated and largely uncharacterized. ncRNAs often originate from genomic regions encompassing enhancers and are generally referred to as enhancer RNAs (eRNAs) (Natoli and Andrau 2012, Li, Notani et al. 2016). eRNAs can be broadly categorized into two types: 2d-eRNAs: Nonpolyadenylated, bidirectional transcripts that are short and heterogeneous in length (Lam, Cho et al. 2013, Li, Notani et al. 2013, Melo, Drost et al. 2013), and 1d-eRNAs: Long, polyadenylated, unidirectional transcripts that often harbor intron-exon structures (Natoli and Andrau 2012, Li, Notani et al. 2016, Trinh, Ummarino et al. 2021). Mounting evidence suggests that eRNAs play a role in long-range gene activation mediated by enhancers (Lam, Cho et al. 2013, Li, Notani et al. 2013, Melo, Drost et al. 2013, Li, Notani et al. 2016). Previously, we demonstrated that the 1d-eRNA *LOUP*, which originates from the URE and extends toward the promoter region, induces PU.1 expression by mediating URE-promoter chromatin looping. This interaction promotes myeloid cell differentiation (Trinh, Ummarino et al. 2021). Yet, outstanding questions remain: How many PU.1-Es generate eRNAs in myeloid cells and are any of them involved in PU.1-mediated regulation of Mono/MΦ development and function? Addressing these questions could yield significant insights into the mechanisms underlying the essential role of PU.1 in Mono/MΦ biology.

In this study, we interrogated the PU.1-mediated transcriptome and cistrome with our comprehensive collection of putative and verified enhancers and discovered that PU.1 and its eRNA *LOUP* are biomolecular components of a novel gene regulatory circuit that mediates the activity of the *LOUP* promoter and a myeloid-specific super-enhancer, promoting their mutual expression. *LOUP* promotes Mono/MΦ differentiation and innate immune functions. Our findings of this myeloid-specific eRNA-TF feed-forward loop (FFL) provide important insights into enhancer regulation underlying Mono/MΦ biology and significant implications for understanding inflammatory diseases and myeloid malignancies.

## METHODS

### Cell lines and Cell Culture

Cell lines (U937, HL-60, K562, HEK293T) were obtained from the American Type Culture Collection (ATCC). The culture conditions for these cell lines were as follows: U937, HL-60, and K562 cells were cultured in RPMI-1640 medium supplemented with 10% (vol/vol) fetal bovine serum (FBS; Cytiva) and 1% penicillin-streptomycin. HEK293T cells were cultured in DMEM medium supplemented with 10% (vol/vol) FBS and 1% penicillin-streptomycin. All cell lines were maintained in incubators at 37°C with 5% (vol/vol) CO2 and humidified atmosphere.

### RNA extraction and RT-qPCR

Cultured cells were centrifuged at 1000 rpm and washed with phosphate-buffered saline (PBS) before isolating RNA. Total RNA was extracted with PureLink™ RNA Mini Kit (Ambion) or Trizol reagent (Invitrogen) following manufacturer’s instruction. Extracted RNA was cleaned for DNA contaminant with RNase-free DNase I (Roche). For RT-qPCR, reverse-transcription was done with the SuperScript® III Reverse Transcriptase (Invitrogen). SYBR® Green Master Mix (Bio-Rad) was used for PCR quantification in Quantstudio5 Real-Time PCR System (Applied Biosystems). Quantification (relative) was performed using the delta CT or standard curve method by comparing Ct values of unknown samples to the standard curve.

### Fluorescence-activated cell sorting and analysis

Surface marker staining and Fluorescence-activated cell sorting (FACS) analysis were performed following our previously described procedures (Mueller, Pabst et al. 2006, Trinh, Ummarino et al. 2021). Briefly, cells were stained with were stained first with Live/Dead ZOMBIE Red (BioLegend, Cat No. 423109), and subsequently with antibodies: anti-CD11b-APC (BD Pharmingen, 340936) and anti-CD15-FITC (Dako, F0830), and anti-CD14-APC-H7 (BD Pharmingen, 641394). Fluorescently labelled cells were analyzed using acoustic focusing technology and Attune NxT cell sorter and data analyzed using FlowJo ver. 10.10 (Tree Star).

### Chemically induced Mono/MΦ and fluorescently labelled beads phagocytosis assay

*LOUP*-depleted and control U937 cells were treated with PMA (50 ng/mL) or solvent control (DMSO) in full RPMI 1640 media (supplemented with 10% FBS, 1% Penicillin-Streptomycin (P/S) and L-glutamine). After 24-hour treatment, cells were transferred to fresh RPMI media without PMA or DMSO and maintained for 72 hours for before fluorescent blue–labelled latex beads (size 2 μm; Sigma; #L0280) were added at 0.005 ng/mL to the culture. After 2-hour incubation, cells were collected and spun down at 300g for 5 minutes at 4°C; pelleted cells were stained with anti-CD11b-APC-Cyanine 7 (BioLegend, 101226) for 30 minutes. Stained cells were fixed with 2% paraformaldehyde for 30 minutes. Fluorescently labelled cells were imaged and analysed using an Amnis ImageStreamX Mark II System at the University of Virginia Flow Cytometry Core Facility. Using 60X objective lens, 10,000 cells were captured from each sample using the INSPIRE® software. Emitted light from blue coloured beads were captured in channel 7 from camera 2 with 420–505 nm filter. Red colour emitted from CD11b were captured in channel 11 with from camera 2 with 660–740 nm filter. All data has been analysed in IDEAS (version 6.2) and FCS Express (version 7).

### Promoter reporter assay

*LOUP* Promoter (wildtype) and its PU.1-binding site mutant version were cloned into pGL3 luciferase reporter plasmid. The assay was performed as described previously (Trinh, Barengo et al. 2011, Trinh, Barengo et al. 2015). In brief, K562 cells co-transfected with plasmid cocktails using lipofectamine 2000 (Invitrogen). These cocktails include firefly luciferase reporter plasmids, *PU.1* cDNA or empty vector, and Renilla luciferase reporter plasmid to normalize transfection efficiency. Luciferase activities were assayed. Luminescence was measured using the Femtomaster FB15 luminometer (Zylux) and the dual-luciferase reporter assay kit (Promega, Madison, WI).

### Enhancer collection

Enhancer coordinates were downloaded from EnhancerAtlas (Gao and Qian 2020), FANTOM5 (Andersson, Gebhard et al. 2014), VistaEnhancer (Visel, Minovitsky et al. 2007). (Visel, Minovitsky et al. 2007), and the human reference genome assembly GRCh38p13 from the Genome Reference Consortium (Nurk, Koren et al. 2022). Enhancer coordinates for nine experimentally validated enhancers that are not present in these databases were annotate manually based on published information (Cockerill, Shannon et al. 1993, Ng, Yokomizo et al. 2010, Levantini, Lee et al. 2011, Guo, Ma et al. 2012, Avellino, Havermans et al. 2016). All coordinates were generated in or converted into GRCh38.

### RNA-seq experiment and analyses

Total RNA with good RNA integrity values (combined 260/280 and 260/230 ≥4) were sequenced ưith NovaSeq PE150 platform (Novogene Corporation Inc). RNA-seq analysis were performed as we described previously (Trinh, Ummarino et al. 2021, Qiu, Vu et al. 2024). Raw sequencing reads (FASTQ files) from in-house and public datasets (GEO) were assessed for read quality by FastQC (Andrews 2016). To remove reads with low-quality trim_galore was used (Krueger 2017). Transcripts per million (TPM) counts were produced using Salmon v1.10.1 with the Ensembl human cDNA catalog GRCh38 (Patro, Duggal et al. 2017). Genes were annotated using Biomart in Ensembl (Hunt, McLaren et al. 2018). Differentially expressed genes (DEGs) are defined as at least two-fold difference between perturbated and control conditions. BRB-ArrayTools was used (Simon, Lam et al. 2007).

### Gene ontology analysis

Database for Annotation, Visualization and Integrated Discovery functional annotation tool (https://davidbioinformatics.nih.gov/) was used for Gene Ontology analysis of biological functions as we have done previously (Trinh, Barengo et al. 2015). Significance of overrepresented biological processes was evaluated based on −log10 of corrected P values using Bonferroni-corrected modified Fisher’s exact test) (Dennis, Sherman et al. 2003).

### Gene Set Enrichment Analysis

Gene Set Enrichment Analysis (GSEA) was conducted using version 4.3.3 of the GSEA software (Subramanian, Tamayo et al. 2005, Trinh, Barengo et al. 2015). The analysis employed the signal-to-noise ratio to score gene sets, which was incorporated into the weighted enrichment score calculation. Statistical significance was determined through 1000 phenotype permutations to generate p-values. The analysis utilized curated gene sets from the Molecular Signatures Database (MSigDB) (Liberzon, Birger et al. 2015) were used.

### ChIP-seq analysis

TF and histone ChIP-seq datasets were obtained from GEO and processed as described in previous studies (Trinh, Ummarino et al. 2021, Qiu, Vu et al. 2024). FastQC was used to assess read quality (Andrews 2016). Low-quality reads were removed using trim_galore when necessary (Krueger 2017). Processed reads were then aligned to the hg38 human genome using the STAR aligner (Dobin, Davis et al. 2013). Coverage maps were generated with bamCoverage, a component of the deepTools suite, using default settings (Ramirez, Ryan et al. 2016). The resulting bigWig files were visualized in the UCSC genome browser. HOMER (v5.1) was employed for peak calling, with peaks showing at least two-fold enrichment over Input control used for annotation (Heinz, Benner et al. 2010). Gene annotation information was retrieved from Ensembl 97 human gene CRCh38.p12 using Biomart (Hunt, McLaren et al. 2018). PU.1-chromatin occupancy profiles were compared with enhancer collections for peak region overlaps using HOMER.

### Molecular docking analysis and molecular interaction visualization

The ETS domain of PU.1 were extracted from Protein Data Bank (PDB: 8E4H) and processed in Pymol Version 2.0 (Schrödinger, LLC). The ETS domain and DNA binding motif on *LOUP* promoter were docked using HDock algorithm (Yan, Zhang et al. 2017).. Full-length human PU.1 protein structure is predicted by AlphaFold (Jumper, Evans et al. 2021). Protein domains with binding structures were visualized and color-coded using Pymol.

## RESULTS

### Genome-wide identification of PU.1-regulated enhancer-associated ncRNAs

We first sought to identify global PU.1 occupancy profile on chromatin in myeloid cells. By analyzing ChIP-seq data of the HL-60 cell line which resemble immature myeloid cells (Gallagher, Collins et al. 1979), we identified 107,797 confident PU.1 ChIP-seq peaks (Figure 1A, Table S1). We next examined the proportion of peaks that are present within gene enhancers. To that end, we compiled a comprehensive collection of enhancers in human. The collection includes well-known online repertories EnhancerAtlas (Gao and Qian 2020), FANTOM5 (Andersson, Gebhard et al. 2014), VistaEnhancer (Visel, Minovitsky et al. 2007), and the human reference genome assembly GRCh38p13 from the Genome Reference Consortium (Nurk, Koren et al. 2022). Nine experimentally validated enhancers that are not present in these databases are also included (Cockerill, Shannon et al. 1993, Ng, Yokomizo et al. 2010, Levantini, Lee et al. 2011, Guo, Ma et al. 2012, Avellino, Havermans et al. 2016) (Figure 1B). By interrogating PU.1 ChIP-seq peaks with the enhancer collections, we revealed 12.7% of PU.1 ChIP-seq peaks being present at 13,096 enhancers (Figures 1A-B). Of note, only a minor fraction of <u>PU.1</u>-bound <u>e</u>nhancers (PU.1-Es) harbor more than one PU.1 ChIP-seq peaks (Figure 1C). Because noncoding RNAs have been increasingly reported to contribute to enhancer activities, we examined the presence of these noncoding genes at the PU.1-Es. By annotating the PU.1-Es with the comprehensive human gene annotation GRCh38.85 from the GENCODE Project, we revealed 7,923 genes that overlap or within 1kilobase from the PU.1-Es. Among these genes, 1,146 (17,05%) are noncoding hereafter referred to as <u>PU.1-E</u>-associated <u>n</u>oncoding <u>g</u>enes (PU.1-ENGs) (Figures 1D and S1A). To identify PU.1-ENGs that are also regulated by PU.1, we first compared RNA-seq transcript profiles between PU.1 knockdown and non-targeting control HL-60 cells thereby identifying 4,758 <u>d</u>ifferentially <u>e</u>xpressed <u>g</u>enes (DEGs) (Figure S1B). Among these, 25.4% are <u>d</u>ifferentially <u>e</u>xpressed <u>n</u>on-coding <u>g</u>enes (DENGs) (Figure S1B). By interrogating PU.1-DENGs with PU.1-ENGs (Figure S1A), we identified 17 NGs that are PU.1-ENGs and PU.1-DENGs (PU.1-E/DENGs) (Figures 1D-E). Of note, most of these NGs are pseudo genes. Intriguingly, we noted the presence of the unidirectional-eRNA (1d-eRNA) *LOUP* that we previously discovered to act as an RNA inducer of *PU.1* (Trinh, Ummarino et al. 2021). The presence of a *PU.1* eRNA that are also induced by PU.1 prompted our further investigation into the role and mechanisms of this biomolecular interplay in epigenetic gene regulation.

**Figure 1.**
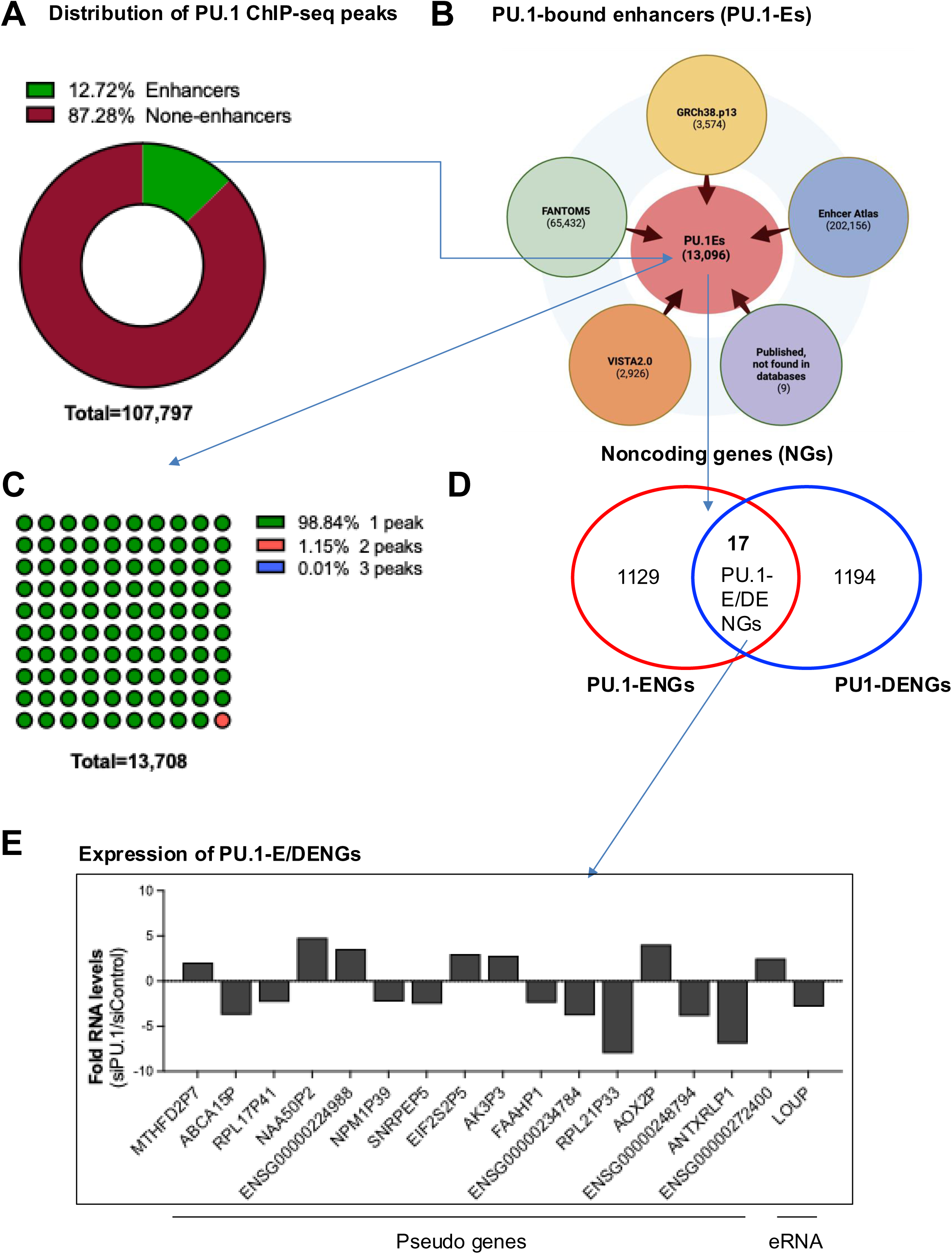
An integrated survey for PU.1-E/DE NGs in HL-60 cells. (A) Pie chart showing distribution of PU.1 ChIP-seq peaks at enhancers and other genomic locations. The peaks were generated from the published dataset GSE32465 that was deposited in Gene Expression Omnibus (GEO). (B) Schematic showing the identification of PU.1-Es from verified and putative enhancer collections. Shown are numbers of enhancers within five collections: FANTOM5, VISTA2.0, Enhancer Atlas, GRCh38.p13, and published literature). (C) Pie chart showing percentage of enhancers exhibiting differential presence of PU.1 ChIP-seq peaks (D) Venn diagram displaying interrogation of PU.1-ENGs (derived from PU.1-Es) and PU.1-DENGs (derived from GSE87055). (E) Bar graph presenting changes in expression levels of PU.1-E-DENGs. Ratio between values PU.1 short-interfering RNA (siRNA) and non-targeting controls (siControl) are shown.

### PU.1 induces eRNA *LOUP* expression within a feed-forward loop (FFL)

To confirm our initial finding of PU.1-mediated *LOUP* expression, we utilized multiple PU.1 perturbational approaches. Knocking down of *PU.1* by siRNAs in the human myeloid cells U937 resulted in robust repression of *LOUP* RNA levels (Figure 2A). In line with this finding, *Pu.1* knockout in murine caused significant reduction of *Loup* in primary murine myeloid cells (Figure 2B). In converse upregulation experiments, enforced *PU.1* expression in K562 cells resulted in strong induction of *LOUP* expression (Figure 2C). Similar results were found in mouse early T cells which are capable of being induced to myeloid lineages (Hosokawa, Ungerback et al. 2018) (Figure 2D). Additionally, releasing *Pu.1* inhibition from the microRNA miR-155 by deleting its recognition element (Lu, Nakagawa et al. 2014) resulted in *Loup* induction in murine splenic B cells (Figure 2E). Furthermore, *PU.1* and *LOUP* expression patterns are highly correlated and predominantly in the myeloid cell lineage (Figure 2F). These data suggest that PU.1 is potent inducer of *LOUP* expression in myeloid cells. Our group and others have previously demonstrated that *LOUP* is also capable of inducing *PU.1* expression (Trinh, Ummarino et al. 2021, Halasz, Malekos et al. 2024). Taken together, our finding indicates that *PU.1* and *LOUP* are protein and eRNA partner of a novel eRNA-TF FFL that positively regulate each other in myeloid cells.

**Figure 2.**
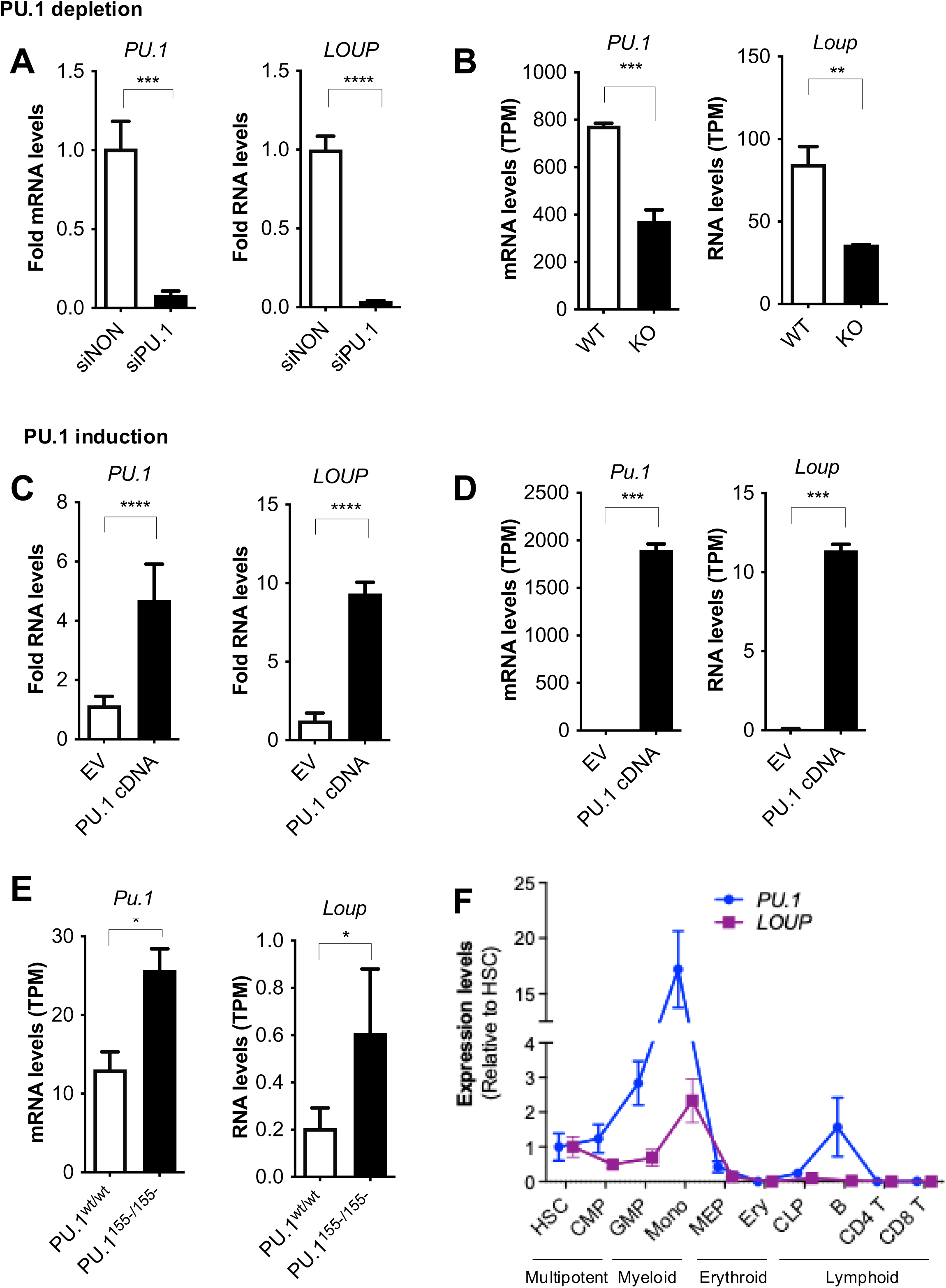
PU.1 induces *LOUP* expression. (A) RT-qPCR expression analysis of *PU.1* (left panel) and of *LOUP* (right panel). U937 cells were transfected with *PU.1*-targeting or non-targeting (control) siRNAs (n=3). (B) Transcript counts of *Pu.1* (left panel) and *Loup* (right panel) in *Pu.1* wildtype and knockout primary murine neutrophils derived from immortalized progenitors (Fischer, Walter et al. 2019). RNA-seq data dataset (GEO: GSE110864, n=3). (C) RT-qPCR expression analysis of *PU.1* (left panel) and of *LOUP* (right panel) in K562 cells transfected with empty vector (EV) or *PU.1* cDNAs (n=3). (D) Transcript counts of *Pu.1* (left panel) and *Loup* (right panel) in the pro-T cell line Scid.adh.2c2 infected with either mock control (EV) or *PU.1* cDNA (Hosokawa, Ungerback et al. 2018), (n=2). Published RNA-seq dataset, GEO: GSE93755. (E) Transcript counts of *Pu.1* (left panel) and *Loup* (right panel) in splenic B cells isolated from wild type and PU.1^[155-/155-]^ knockin mice (Lu, Nakagawa et al. 2014) (GEO: GSE162526, n>=4). (F) Transcript profile of *PU.1* and *LOUP* in normal blood cell populations. Shown are RNA-seq transcript counts (published dataset GEO: GSE74246). Cell populations include Hematopoietic Stem Cells (HSC), Common Myeloid Progenitors (CMP), Granulocyte-Myeloid Progenitors (GMP), Myeloid (represented by monocytic cells), Myeloid-Erythroid Progenitors (MEP), Erythroid (Ery), Common Lymphoid Progenitors (CLP), B cells, and T cells (include T_CD4+_ and T_CD8+_). Error bars indicate SD. Error bars indicate SD, \**p* < 0.05, \*\**p* < 0.01, \*\*\**p* < 0.001, \*\*\*\**p* < 0.0001.

### PU.1 induces *LOUP* expression by occupying cis-regulatory elements and regulates promoter activity

The human *LOUP* locus spans from the 5’ end of the ∼8 kilobase PCREC (Figure 3A)(Qiu, Vu et al. 2024). Chromatin at the PCREC is accessible, corresponding to the presence of DNase I hypersensitive sites. PCREC is also enriched with histone H3 lysine 27 acetylation (H3K27ac), the active enhancer histone mark (Creyghton, Cheng et al. 2010) as well as histone H3 lysine 4 mono-methylation (H3K4me1), the histone marks associated with the existence of enhancers and gene transcription (Pekowska, Benoukraf et al. 2011) (Figure 3A, epigenetic panels). Thus, *LOUP* locus is located within the PCREC that exhibits features of myeloid-specific enhancers. To understand the molecular mechanism underlying PU.1-mediated regulation of the PCREC within the PU.1-*LOUP* FFL, we examined PU.1 occupancy at the chromatin spanning the *LOUP* locus. PU.1 occupies several conserved PCREs in myeloid cells, whereas its occupancy is limited to fewer locations in B and erythroid cell lineages (Figure 3A, PU.1 ChIP-seq and conservation panels). Within the PCREC, the URE, which overlap with the core promoter region and the first exon of *LOUP* (Figure 1B), carries two PU.1 ChIP-seq peaks. Of note, the first peak is myeloid specific and located at the <u>h</u>omology region 1 (H1) within the URE (Korczmar, Bookstaver et al. 2024) corresponding to the core promoter region of *LOUP* (Figures 3A-B). Whereas the second peak is commonly occupied by PU.1 in multiple cell types and located at the H2 URE (Korczmar, Bookstaver et al. 2024) (Figures 3A-B).

**Figure 3.**
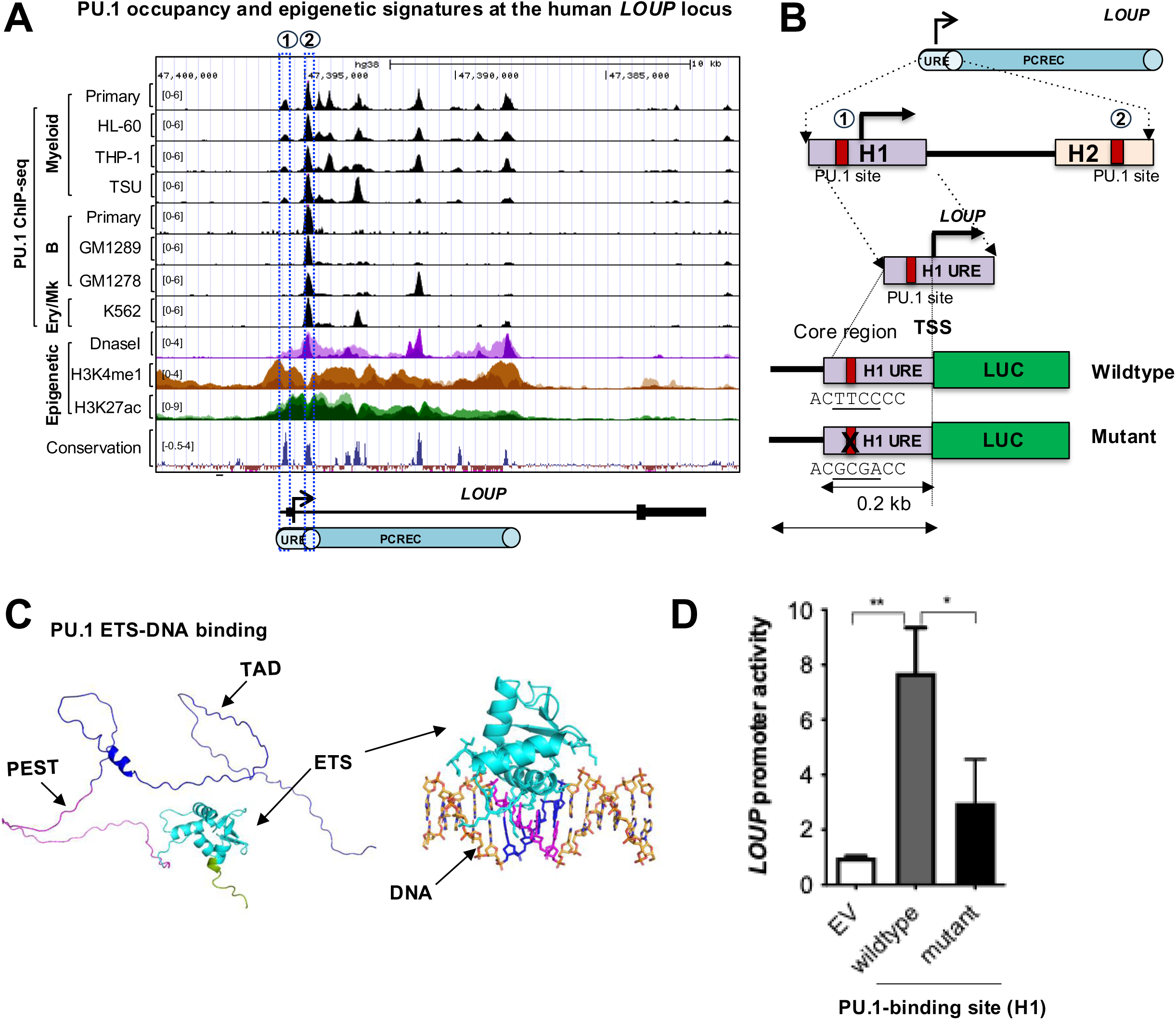
PU.1 occupy the *LOUP* locus and induce it promoter activity. A) Gene track view of the genomic region containing the human *LOUP* locus. Top panels include PU.1 ChIP-seq tracks of primary and blood cell lines representing myeloid (primary cells, HL-60, THP-1 and TSU cell lines) B (primary cells, GM1289 and GM1278 cell lines), and erythroid/megakaryocytic cell lineages (K562 cell line). Middle panels include epigenetic tracks of overlay DNAse-seq, and ChIP-seq for enhancer marks (H3K4me1 and H3K27ac) from myeloid cell lines (More details about the samples are included in the Method section). Bottom panels include a PhyloP of 100 vertebrates Basewise Conservation and a schematic diagram (underneath) showing the genomic location of the *LOUP* locus with the corresponding PCREC. The location of the major PCRE, URE is shown. Circles with numbers 1 and 2 and dotted-line boxes denote two PU.1 ChIP-seq peaks within the URE. B) Schematic diagram of the *LOUP* promoter reporter constructs. Relative position of the *LOUP* core promoter and corresponding PCREC are shown. Within the PCREC, the URE with H1 and H2 regions are illustrated. Red boxes denote PU.1 binding motifs (without “X”: wildtype, with “X”: point mutations abrogating the PU.1 binding motif (mutant)). Underneath are nucleotide sequences for the wildtype and mutant motifs. C) PU.1 domain structure and molecular simulation showing interaction between the PU.1 ETS domain and *LOUP* promoter. Left panel: AlphaFold prediction of full-length human PU.1 protein structure. Arrows point to PU.1 domains which are color-coded (ETS in cyan, PEST in magentas, and TAD in blue). Right panel: Molecular simulation showing interaction between the PU.1 ETS domain (PDB: 8E4H) and the PU.1 binding motif at the *LOUP* promoter region. ETS domain is colored as cyan with DNA interacting interface shown as sticks. DNA is colored as orange. PU.1 core binding motif (underlined figure B) are highlighted in magenta (sense strand) and blue (anti-sense strand). D) Reporter assay in K562 cells transfected with PU.1 cDNA together with wildtype or mutant *LOUP* reporter constructs (shown in (B)), respectively. Fold changes in compared to empty vector (EV) control are shown.

Because the URE contains two major *LOUP* elements with PU.1 occupancy (the core promoter and first exon) (Trinh, Ummarino et al. 2021, Korczmar, Bookstaver et al. 2024) and have been shown to play a critical role in inducing *PU.1* expression (Trinh, Ummarino et al. 2021, Korczmar, Bookstaver et al. 2024), we focused our analyses on this PCRE. In examining the DNA sequences of these elements, we identified a PU.1 consensus binding motif within the PU.1 ChIP-seq peak located within the core *LOUP* promoter region (Figure 3B). The human PU.1 protein with 264 amino acids contain three functional domains, the N-terminal transactivation domain (TAD), a centrally located PEST domain with abundances of proline (P), glutamic acid (E), serine (S) and threonine (T) residues, and the C-terminal ETS domain that binds DNA (Figure 3C, left panel). In line with previous reports that PU.1 binds its consensus DNA motif via the ETS domain (Klemsz and Maki 1996, Petrovick, Hiebert et al. 1998), our molecular simulation analysis visualize interaction between this domain to the binding motif at the core *LOUP* promoter region (Figure 3C, right panel). By performing promoter reporter assay, we showed that PU.1 induces *LOUP* promoter activity. Additionally, point mutations abrogating PU.1 binding motif caused a significant reduction in *LOUP* promoter activity (Figures 3B and 3D). These data indicate that PU.1 induces *LOUP* promoter through binding to the PU.1-consensus motif within the URE.

### *LOUP* induces Mono/MΦ differentiation

We previously demonstrated that *LOUP* induces myeloid differentiation (Trinh, Ummarino et al. 2021). However, the role of this eRNA in Mono/MΦ differentiation and their cellular functions has not been closely examined. To elucidate this, we depleted *LOUP* in CD34^+^ hematopoietic stem/progenitor cells (HSPCs) using short hairpin RNAs (shRNAs) and cultured the cells with myeloid differentiation-promoting cytokines (Figure 4A). Knocking down *LOUP* (Figure 4B) led to a decrease in the expression of the mono/macrophage marker CD14 (Figures 4C, upper panel, and S2A). As expected, expression of the general marker for human myeloid cell differentiation CD15 is also reduced (Figures 4C, lower panel). We further depleted *LOUP* by shRNA in the pro-monocytic cell line U937 (Sundstrom and Nilsson 1976) (Figure 4D). This cell line can be chemically induced to undergo Mono/MΦ differentiation by Phorbol 12-myristate 13-acetate (PMA) (Figure 4E) as evidenced by changes in cell morphology from floating and ball-like to adherent and spindle-like (Figure S2B), as well as increases in general myeloid and Mono/MΦ markers *CD11B* and *CD14* (Figures S4C-D). *LOUP* depletion counteracts the effect of PMA-induced expression of the mono/MΦ marker *CD14* (Figures 4F and S2E). Similar finding was observed when *LOUP* was depleted by CRISPR/Cas9 technology (Figures 4G and S2E-F). We further examined global changes in gene expression that are mediated by *LOUP* using Gene Set Enrichment Analysis (GSEA) (Subramanian, Tamayo et al. 2005, Trinh, Barengo et al. 2015). *LOUP* depletion resulted in gene expression programs that were inversely correlated with two gene sets that signify differentiation of monocytes (Figure 4H) and MΦ (Figure 4I), respectively, suggesting that the *LOUP*-mediated transcript profile is associated with Mono/MΦ differentiation. Taken together, these results demonstrate that *LOUP* mediate the Mono/MΦ transcript profile thereby promoting Mono/MΦ differentiation.

**Figure 4.**
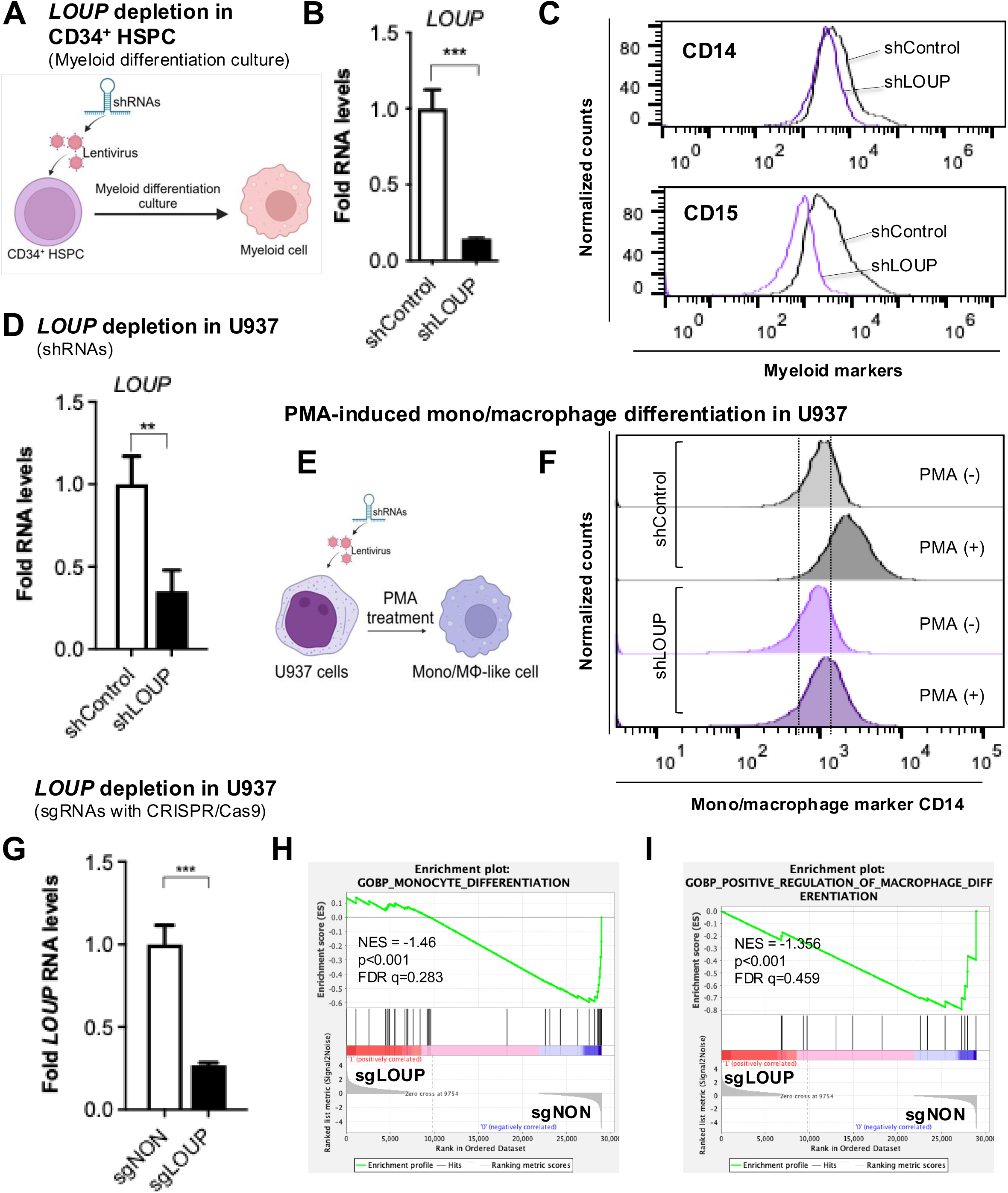
*LOUP* induces Mono/MΦ differentiation. A) Schematic illustrating *ex vivo* myeloid differentiation. Cord blood CD34^+^ HSPC cells were transduced with shRNAs targeting *LOUP* (shLOUP) and non-targeting control shRNA (shControl). Transduced cells were cultured in media containing myeloid differentiation-promoting cytokines including granulocyte-macrophage colony-stimulating factor (GM-CSF), granulocyte colony-stimulating factor (G-CSF), and Interleukin 3 (IL-3) for 12 days. B) RT-qPCR analysis of *LOUP* expression after *ex vivo* myeloid differentiation as described (A). C) Representative FACS results of myeloid marker expression CD14 and CD15) as described (A). D) RT-qPCR analysis of *LOUP* expression in U937 cells that were transfected by lentiviral transduction with either shControl shRNA or shLOUP shRNA. E) Schematic illustrating PMA-induced Mono/MΦ differentiation. Upon transduction with shRNAs targeting *LOUP* (shLOUP) or non-targeting control shRNA (shControl), pro-monocytic cell line U937 cells were culture in 10nM PMA for 72h hours. F) Representative FACS results of myeloid marker expression CD14 as described (E). G) RT-qPCR analysis of *LOUP* expression in *LOUP*-depleted U937 (sgLOUP) and control (sgControl) CRISPR/Cas9) via CRISPR/Cas9. H-I) GSEA analyses for global gene expression changes mediated by *LOUP* in U937 cells using sgLOUP and sgControl cells. Enrichment of monocytic gene sets that signify differentiation of monocytes (H) and MΦ (I). Indicated are normalized enrichment scores (NES), significance values and false discovery rates (FDR).

### *LOUP* promotes major mono/MΦ defense mechanisms against infections

We extended our transcriptomic analysis of global gene expression changes to identify biological processes mediated by *LOUP* using GSEA with the Molecular Signatures Database (MSigDB), one of the most comprehensive collection of gene expression signatures (Liberzon, Birger et al. 2015). The analyses revealed top enriched gene sets that are related to innate immune responses mediated by mono/MΦ (Figure 5A). In parallel, we integrated transcriptome comparative analyses with Gene Ontology (GO) analysis (Dennis, Sherman et al. 2003, Trinh, Barengo et al. 2015) to identify top biological processes mediated by *LOUP.* Specifically, by comparing between *LOUP*-depleted (sgLOUP) and non-targeting control (sgControl) transcript profiles, we identified a repertoire of *LOUP-*mediated DEGs (Figure S3A). By evaluating genes that are induced by *LOUP* (i.e., genes that are downregulated upon *LOUP* depletion (Figure S3A, left)) using GO analysis, we identified top biological processes associated with *LOUP* expression (Figure 5B). In line with GSEA finding, these biological processes are related to innate mono/MΦ defense mechanisms (Figure 5B). Specifically, phagocytosis in which mono/MΦ engulfs pathogens into phagosomes (Uribe-Querol and Rosales 2020) is among the top enriched biological processes (Figure 5B). To further elucidate this important cellular feature, we performed engulfment assays using blue-colored beads coupled with imaging flow cytometry which combines the quantitative capabilities of flow cytometry with the visual confirmation provided by imaging (Headland, Jones et al. 2014). As expected, PMA-induced U937 mono/MΦ show increased cellular uptake of fluorescent-labeled particles (Figures 5C and S3B). However, *LOUP* depletion dampened this effect (Figures 5C-D) indicating that *LOUP* confers Mono/MΦ phagocytic activities. Within the phagosomes, antimicrobial activities such as production of reactive oxygen and nitrogen species (superoxide generation, nitric-oxide synthase) is eminent (Piacenza, Trujillo et al. 2019). In line with this, gene sets and GO terms linked to antimicrobial activities are enriched with *LOUP* expression (Figures 5A-B). Moreover, as part of innate immune defenses, infected Mono/MΦ often produce cytokines to attract and activate other immune cells (Arango Duque and Descoteaux 2014). Accordingly, chemokine production is also induced by *LOUP* (Figure 5B, GO:0032722). Taken together, our results indicates that *LOUP* promotes major mono/MΦ defense mechanisms against infections including phagocytosis, antimicrobial activities and chemoattractant cytokine production.

**Figure 5.**
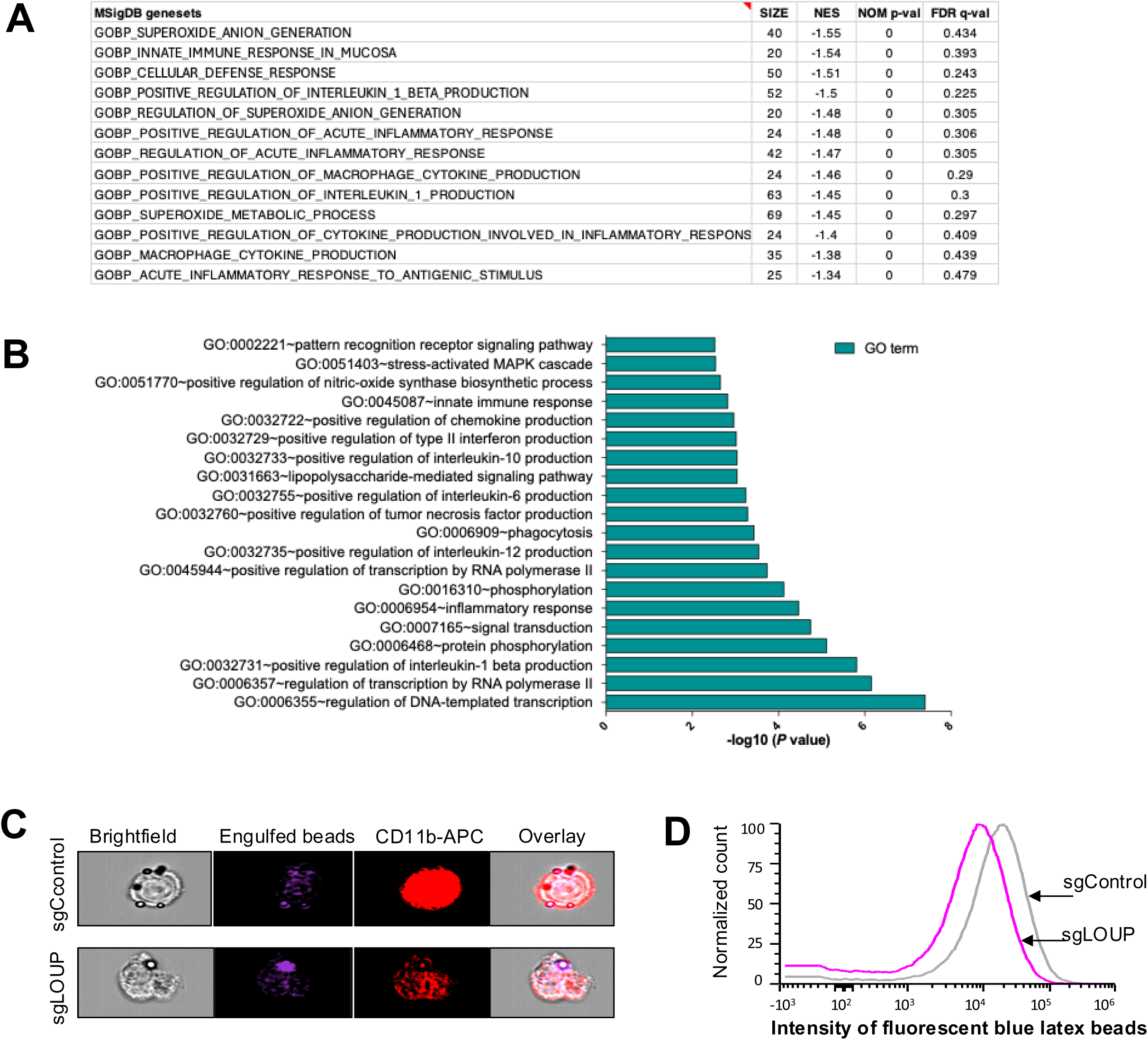
*LOUP* induces Mono/MΦ phagocytic activities. A) GSEA analysis with a collection of gene expression signatures from MSigDB (Liberzon, Birger et al. 2015) for global gene expression changes in U937 cells using sgLOUP and sgControl cells. Shown are top enriched gene sets with associated NES, significance values, and FDR. B) Gene Ontology analysis showing top GO terms induced by *LOUP* in the pro-Mono cell line U937. - log10 of p-value are shown. C-D) Imaging flow cytometry analysis of *LOUP*-depleted (sgLOUP) vs control (sgControl). Shown are representative images of cells with brightfield, immune fluorescence, Engulfed beads, CD11b-APC, and overlay (C). Representative FACS results of fluorescent intensity for engulfed fluorescent blue latex beads as described (D).

In summary, we conducted a systematic genome-wide identification of PU.1-regulated enhancer-associated ncRNAs by interrogating the PU.1-mediated transcriptome and cistrome with our comprehensive collection of putative and verified enhancers. In doing so, we discovered that PU.1 and its eRNA *LOUP* are biomolecular components of a novel gene regulatory circuit that mediates the activity of the *LOUP* promoter and a myeloid-specific super-enhancer, promoting their mutual expression, mono/MΦ differentiation and innate defense mechanisms (Figure 6).

**Figure 6.**
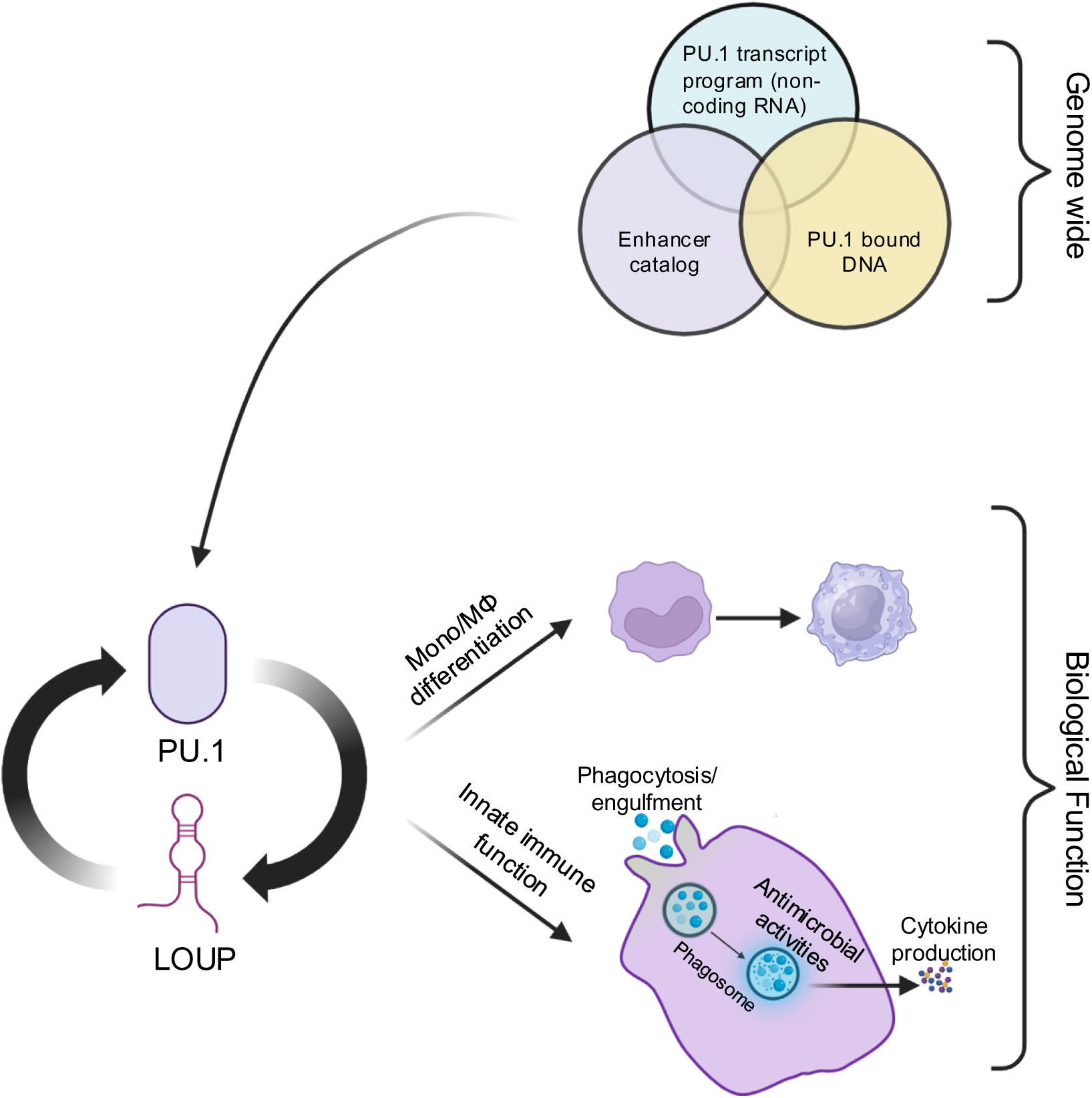
A schematic explaining the role and mechanism of the *LOUP*-*PU.1* FFL.

## DISCUSSION

In this study, we discovered that PU.1 coordinates with *LOUP* at the chromatin level to establish a myeloid-specific eRNA-TF FFL that promotes Mono/MΦ differentiation and enhances innate immune functions. Our study highlights several key findings: First, we demonstrated that PU.1 acts as a potent TF inducer of the 1d-eRNA *LOUP*, which we previously identified as an RNA regulator of PU.1 (Trinh, Ummarino et al. 2021). PU.1 drives *LOUP* expression by binding to a specific DNA motif within the PCREC that exhibits features of a myeloid-specific super-enhancer. The PCREC encompasses the *LOUP* promoter and the URE. Second, we found that *LOUP* plays a pivotal role in promoting Mono/MΦ differentiation by orchestrating the transcriptomic profile characteristic of these cells. Third, *LOUP* promotes critical Mono/MΦ defense mechanisms, including phagocytosis, antimicrobial activity, and the production of chemoattractant cytokines. These functions underscore its essential role in innate immune responses. Collectively, our findings provide valuable insights into how the PU.1/*LOUP* regulatory circuit governs Mono/MΦ differentiation and innate immune function. The identification of this novel eRNA-TF FFL may have significant implications for advancing our understanding of inflammatory diseases and myeloid malignancies.

Our discovery of the FFL mediated by the myeloid-specific eRNA *LOUP* and the TF PU.1 provides critical mechanistic insights into cell-type-specific enhancer regulation. Prior studies, including our own, have shown that PU.1 autoregulates its expression (Staber, Zhang et al. 2013, Korczmar, Bookstaver et al. 2024, Qiu 2024). However, the mechanism underlying this process remained elusive. Unexpectedly, we found that PU.1 achieves this by strongly inducing expression of *LOUP* by activating *LOUP* promoter. Previously, we demonstrated that *LOUP* coordinates with TFs to mediate enhancer-induced PU.1 expression (Trinh, Ummarino et al. 2021). This suggests that *LOUP* and PU.1 act as partner regulatory biomolecules within an enhancer-mediated FFL that mutually reinforces their expression in myeloid cells, thereby driving Mono/MΦ differentiation and innate immune function. To our knowledge, this is the first description of a myeloid-specific FFL in which the target gene of the eRNA also functions as a potent inducer of eRNAs via modulating enhancer activity. We propose that such gene circuits are pivotal for driving robust target gene expression providing important mechanistic insights into gene-specific enhancer-promoter interactions and cell-type-specific transcriptional regulation.

There are several inherent experimental challenges associated with studying ncRNAs, especially eRNAs. The first challenge arises from the biological characteristics of noncoding RNAs, such as their low expression levels (Zhao, Teng et al. 2020). To prevent false-positive detection caused by quantifying contaminating genomic DNA or unrelated RNAs, we designed primers targeting exon junctions of *LOUP* and implemented a depletion step to remove contaminated genomic DNA prior to RT-qPCR. These precautions were taken to ensure accurate detection and quantification. Another significant challenge stems from the overlap in genomic location between eRNAs and enhancers, which introduces technical limitations in distinguishing the activity of eRNAs from that of the enhancers themselves. Directly targeting eRNAs at the genomic level often risks inadvertently disrupting enhancer function. For example, recruiting dCas9-KRAB inhibitors to target *LOUP* at the homology region 1 of the URE (PU.1 enhancer) could directly interfere with enhancer activity (Ebralidze, Guibal et al. 2008, Halasz, Malekos et al. 2024).

To address this issue, we applied multiple approaches and avoided targeting the genomic region containing the enhancer. For instance, our CRISPR knockout strategies were specifically designed to avoid directly targeting the URE. Additionally, we employed shRNA-mediated knockdown of *LOUP*, which targets *LOUP* at the RNA level. By utilizing complementary approaches while accounting for low RNA expression levels, we strive to overcome the inherent technical hurdles associated with studying noncoding RNAs like eRNAs.

Our findings indicate the important role of the PU.1/*LOUP* FFL in normal and malignant myelopoiesis. PU.1 has long been known to be essential for myeloid development and is required for terminal Mono/MΦ differentiation (Tenen 2003). PU.1 down regulation impairs myeloid cell differentiation, resulting in the development of acute myeloid leukemia (AML) (Cook, McCaw et al. 2004, Rosenbauer, Wagner et al. 2004). On the other hand, we previously found that *LOUP* induces PU.1 expression and promotes myeloid development. *LOUP* also exhibits AML inhibiting functions and is repressed by oncogenic fusion proteins AML (Trinh, Ummarino et al. 2021). Here, we demonstrate that PU.1 is a powerful TF inducer of *LOUP*, and that *LOUP* induces Mono/MΦ differentiation. Taken together, our findings, combined with the aforementioned reports, point to the role of the PU.1/*LOUP* FFL in myeloid differentiation in general and Mono/MΦ differentiation specifically. As this gene regulatory circuit is downregulated in AML, it could represent an Achilles’ heel in AML, potentially to be modulated therapeutically.

Our findings highlight the potential role of the PU.1/*LOUP* FFL in innate immune functions of Mono/MΦ. In particular, *LOUP* promotes major Mono/MΦ defense mechanisms against infections, including phagocytosis, antimicrobial activities, and chemoattractant cytokine production. A recent study suggests that *LOUP* may produce small peptides, suggesting additional molecular functions of this eRNA (Halasz, Malekos et al. 2024). On the other hand, PU.1 governs innate immune responses, playing a role in pathological conditions involving Mono/MΦ, such as inflammation related to asthma and allergies (Karpurapu, Wang et al. 2011, Qian, Deng et al. 2015, Fischer, Walter et al. 2019). Because PU.1 and *LOUP* induce each other within the PU.1/*LOUP* FFL, disentangling the contributions of individual components within the FFL could be challenging due to their mutual regulatory nature. Nevertheless, the effect of each component could apparently be attributed collectively to the PU.1/*LOUP* FFL’s output. Further research will identify upstream regulator as well as downstream cytokine effectors of this FFL. These insights could further illuminate the role and mechanism of this regulatory circuit in in development and immune functions of Mono/MΦ pave the way for innovative treatments aimed at restoring immune homeostasis and enhancing pathogen clearance.

In summary, we uncovered a fascinating interplay between the TF PU.1 and its eRNA *LOUP*, which form a myeloid-specific eRNA-TF FFL. This regulatory mechanism operates through chromatin regulation, enabling PU.1 and *LOUP* to induce each other’s expression. Our findings shed light on the intricate processes of cell-type specific and enhancer regulation in Mono/MΦ, providing valuable insights into enhancer function. Importantly, we discovered that *LOUP*, as a component of this FFL, plays a crucial role in mediating Mono/MΦ differentiation and innate immune function. This revelation suggests that this regulatory hub may have far-reaching implications for our understanding of inflammatory diseases and myeloid malignancies, potentially opening new avenues for research and therapeutic interventions.

## Supporting information

None

## ACKNOWLEDGEMENTS

We thank UVA Research Computing staff, Satish Tiwari, Yanzhou Zhang, Huong Trinh, Danielle Llaneza, Yaning Li, Jenna Sleiman, Emilia Korczmar, Anna Bookstaver, and members of the Trinh laboratory for assistance and helpful suggestions. Figure 1B, 4A, 4E, 6 were created via BioRender.

---

**Fig S1.**
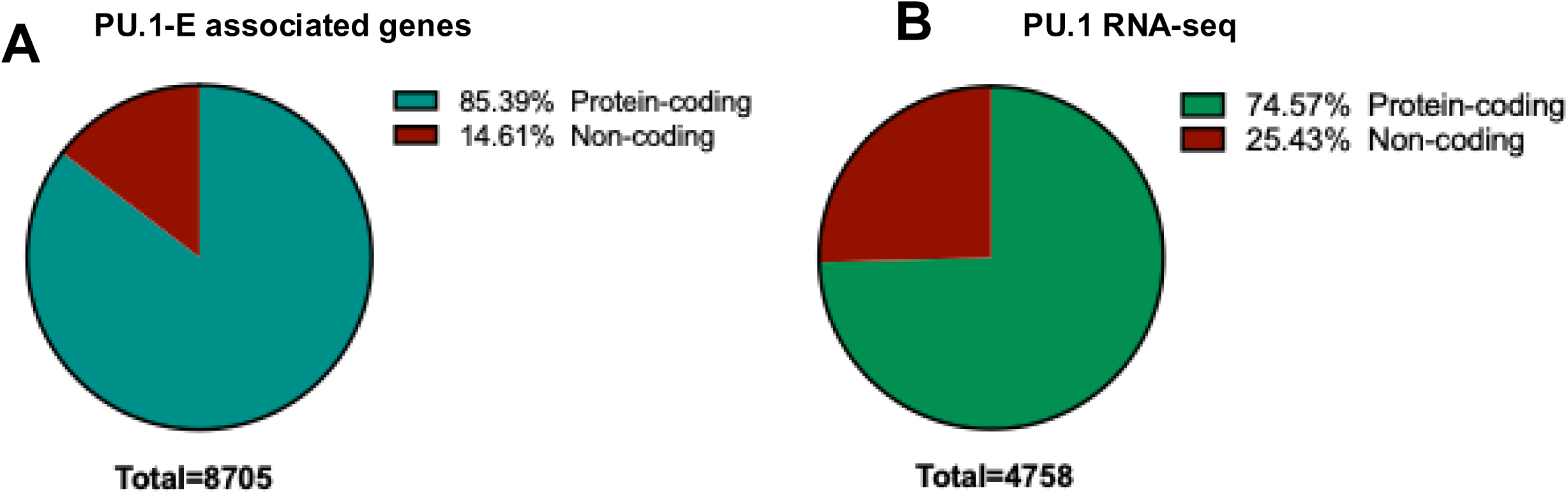

**Fig S2.**
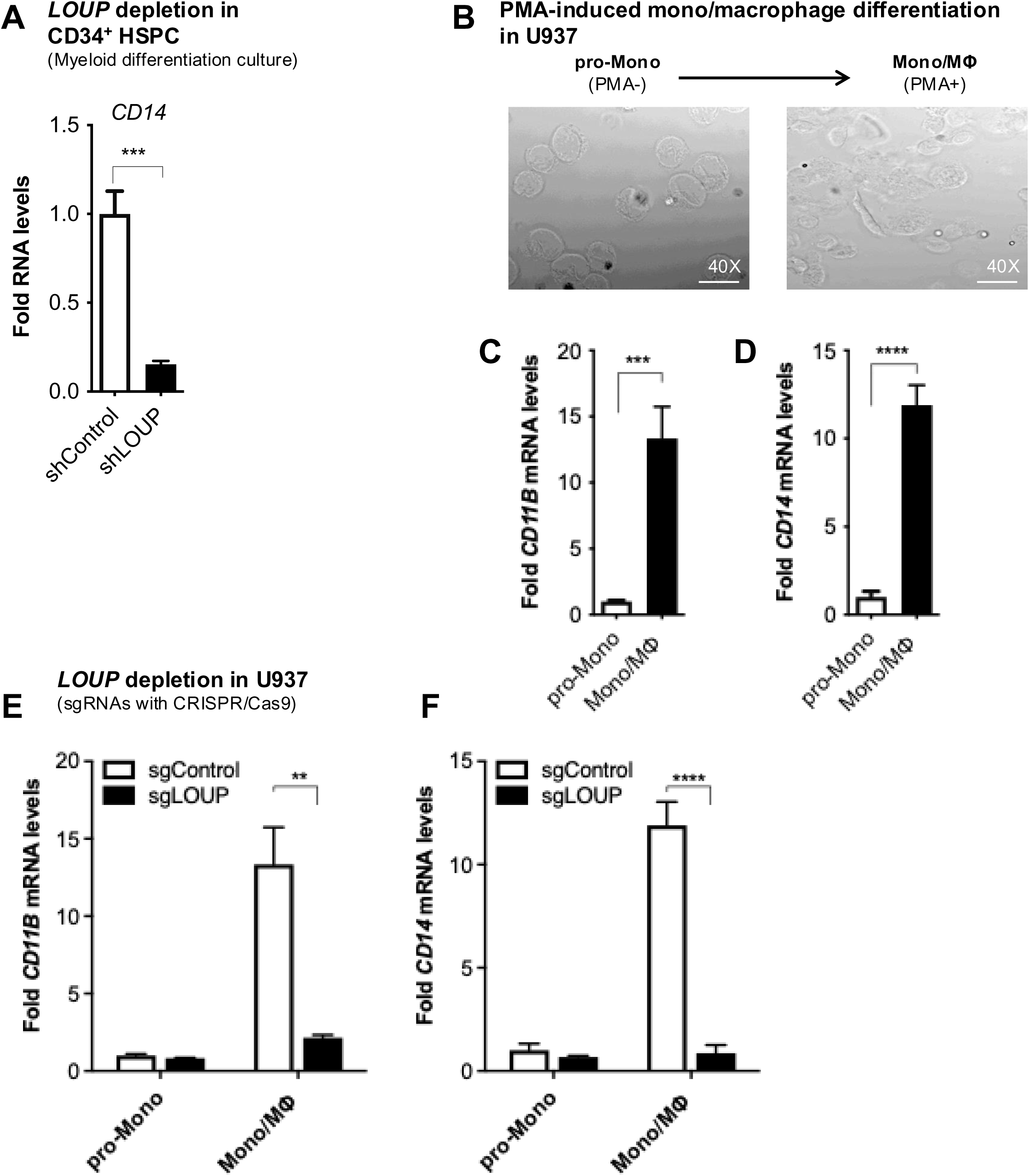

**Fig S3.**
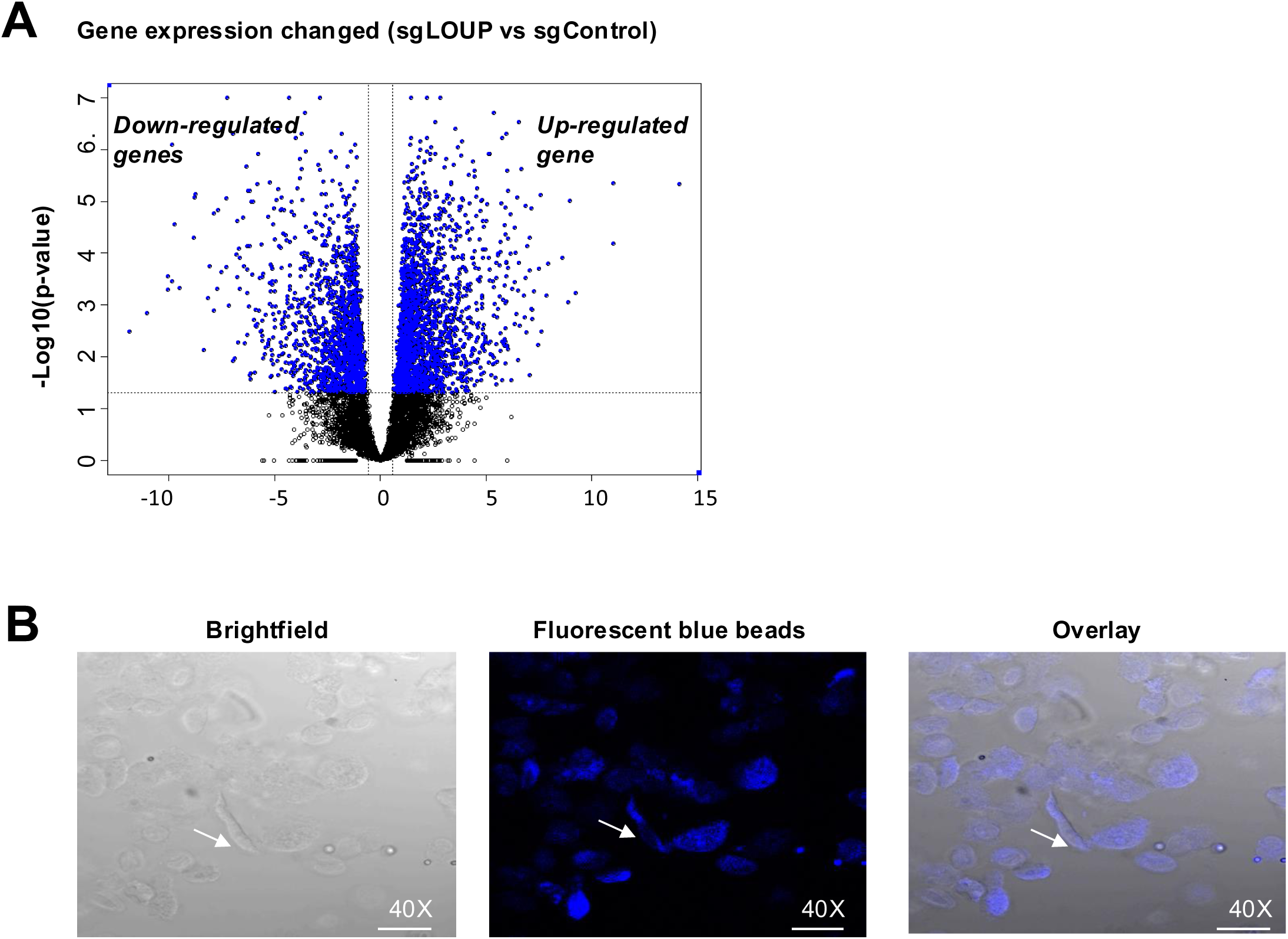

## REFERENCES

Andersson, R., C. Gebhard, I. Miguel-Escalada, I. Hoof, J. Bornholdt, M. Boyd, Y. Chen, X. Zhao, C. Schmidl, T. Suzuki, E. Ntini, E. Arner, E. Valen, K. Li, L. Schwarzfischer, D. Glatz, J. Raithel, B. Lilje, N. Rapin, F. O. Bagger, M. Jorgensen, P. R. Andersen, N. Bertin, O. Rackham, A. M. Burroughs, J. K. Baillie, Y. Ishizu, Y. Shimizu, E. Furuhata, S. Maeda, Y. Negishi, C. J. Mungall, T. F. Meehan, T. Lassmann, M. Itoh, H. Kawaji, N. Kondo, J. Kawai, A. Lennartsson, C. O. Daub, P. Heutink, D. A. Hume, T. H. Jensen, H. Suzuki, Y. Hayashizaki, F. Muller, A. R. R. Forrest, P. Carninci, M. Rehli and A. Sandelin (2014). "An atlas of active enhancers across human cell types and tissues." Nature 507(7493): 455–461.

Andrews, S. (2016). "FastQC: a Quality Control Tool for High Throughput Sequence Data." Babraham Bioinformatics version 0.11.5, from https://www.bioinformatics.babraham.ac.uk/projects/fastqc/.

Arango Duque, G. and A. Descoteaux (2014). "Macrophage cytokines: involvement in immunity and infectious diseases." Front Immunol 5: 491.

Avellino, R., M. Havermans, C. Erpelinck, M. A. Sanders, R. Hoogenboezem, H. J. van de Werken, E. Rombouts, K. van Lom, P. M. van Strien, C. Gebhard, M. Rehli, J. Pimanda, D. Beck, S. Erkeland, T. Kuiken, H. de Looper, S. Groschel, I. Touw, E. Bindels and R. Delwel (2016). "An autonomous CEBPA enhancer specific for myeloid-lineage priming and neutrophilic differentiation." Blood 127(24): 2991–3003.

Barozzi, I., M. Simonatto, S. Bonifacio, L. Yang, R. Rohs, S. Ghisletti and G. Natoli (2014). "Coregulation of transcription factor binding and nucleosome occupancy through DNA features of mammalian enhancers." Mol Cell 54(5): 844–857.

Chen, H., D. Ray-Gallet, P. Zhang, C. J. Hetherington, D. A. Gonzalez, D. E. Zhang, F. Moreau-Gachelin and D. G. Tenen (1995). "PU.1 (Spi-1) autoregulates its expression in myeloid cells." Oncogene 11(8): 1549–1560.

Chen, H. M., P. Zhang, M. T. Voso, S. Hohaus, D. A. Gonzalez, C. K. Glass, D. E. Zhang and D. G. Tenen (1995). "Neutrophils and monocytes express high levels of PU.1 (Spi-1) but not Spi-B." Blood 85(10): 2918–2928.

Cockerill, P. N., M. F. Shannon, A. G. Bert, G. R. Ryan and M. A. Vadas (1993). "The granulocyte-macrophage colony-stimulating factor/interleukin 3 locus is regulated by an inducible cyclosporin A-sensitive enhancer." Proc Natl Acad Sci U S A 90(6): 2466–2470.

Cook, W. D., B. J. McCaw, C. Herring, D. L. John, S. J. Foote, S. L. Nutt and J. M. Adams (2004). "PU.1 is a suppressor of myeloid leukemia, inactivated in mice by gene deletion and mutation of its DNA binding domain." Blood 104(12): 3437–3444.

Cooper, S., H. Guo and A. D. Friedman (2015). "The +37 kb Cebpa Enhancer Is Critical for Cebpa Myeloid Gene Expression and Contains Functional Sites that Bind SCL, GATA2, C/EBPalpha, PU.1, and Additional Ets Factors." PLoS One 10(5): e0126385.

Costa, F. F. (2008). "Non-coding RNAs, epigenetics and complexity." Gene 410(1): 9–17.

Creyghton, M. P., A. W. Cheng, G. G. Welstead, T. Kooistra, B. W. Carey, E. J. Steine, J. Hanna, M. A. Lodato, G. M. Frampton, P. A. Sharp, L. A. Boyer, R. A. Young and R. Jaenisch (2010). "Histone H3K27ac separates active from poised enhancers and predicts developmental state." Proc Natl Acad Sci U S A 107(50): 21931–21936.

De Kleer, I., F. Willems, B. Lambrecht and S. Goriely (2014). "Ontogeny of myeloid cells." Front Immunol 5: 423.

Dennis, G., Jr., B. T. Sherman, D. A. Hosack, J. Yang, W. Gao, H. C. Lane and R. A. Lempicki (2003). "DAVID: Database for Annotation, Visualization, and Integrated Discovery." Genome Biol 4(5): P3.

Dobin, A., C. A. Davis, F. Schlesinger, J. Drenkow, C. Zaleski, S. Jha, P. Batut, M. Chaisson and T. R. Gingeras (2013). "STAR: ultrafast universal RNA-seq aligner." Bioinformatics 29(1): 15–21.

Ebralidze, A. K., F. C. Guibal, U. Steidl, P. Zhang, S. Lee, B. Bartholdy, M. A. Jorda, V. Petkova, F. Rosenbauer, G. Huang, T. Dayaram, J. Klupp, K. B. O’Brien, B. Will, M. Hoogenkamp, K. L. Borden, C. Bonifer and D. G. Tenen (2008). "PU.1 expression is modulated by the balance of functional sense and antisense RNAs regulated by a shared cis-regulatory element." Genes Dev 22(15): 2085–2092.

Faust, N., C. Bonifer and A. E. Sippel (1999). "Differential activity of the −2.7 kb chicken lysozyme enhancer in macrophages of different ontogenic origins is regulated by C/EBP and PU.1 transcription factors." DNA Cell Biol 18(8): 631–642.

Fischer, J., C. Walter, A. Tonges, H. Aleth, M. J. C. Jordao, M. Leddin, V. Groning, T. Erdmann, G. Lenz, J. Roth, T. Vogl, M. Prinz, M. Dugas, I. D. Jacobsen and F. Rosenbauer (2019). "Safeguard function of PU.1 shapes the inflammatory epigenome of neutrophils." Nat Immunol 20(5): 546–558.

Gallagher, R., S. Collins, J. Trujillo, K. McCredie, M. Ahearn, S. Tsai, R. Metzgar, G. Aulakh, R. Ting, F. Ruscetti and R. Gallo (1979). "Characterization of the continuous, differentiating myeloid cell line (HL-60) from a patient with acute promyelocytic leukemia." Blood 54(3): 713–733.

Gao, T. and J. Qian (2020). "EnhancerAtlas 2.0: an updated resource with enhancer annotation in 586 tissue/cell types across nine species." Nucleic Acids Res 48(D1): D58–D64.

Ginhoux, F. and S. Jung (2014). "Monocytes and macrophages: developmental pathways and tissue homeostasis." Nat Rev Immunol 14(6): 392–404.

Groen, J. N., D. Capraro and K. V. Morris (2014). "The emerging role of pseudogene expressed non-coding RNAs in cellular functions." Int J Biochem Cell Biol 54: 350–355.

Guo, H., O. Ma, N. A. Speck and A. D. Friedman (2012). "Runx1 deletion or dominant inhibition reduces Cebpa transcription via conserved promoter and distal enhancer sites to favor monopoiesis over granulopoiesis." Blood 119(19): 4408–4418.

Ha, S. D., W. Cho, R. P. DeKoter and S. O. Kim (2019). "The transcription factor PU.1 mediates enhancer-promoter looping that is required for IL-1beta eRNA and mRNA transcription in mouse melanoma and macrophage cell lines." J Biol Chem 294(46): 17487–17500.

Halasz, H., E. Malekos, S. Covarrubias, S. Yitiz, C. Montano, L. Sudek, S. Katzman, S. J. Liu, M. A. Horlbeck, L. Namvar, J. S. Weissman and S. Carpenter (2024). "CRISPRi screens identify the lncRNA, LOUP, as a multifunctional locus regulating macrophage differentiation and inflammatory signaling." Proc Natl Acad Sci U S A 121(22): e2322524121.

Headland, S. E., H. R. Jones, A. S. D’Sa, M. Perretti and L. V. Norling (2014). "Cutting-edge analysis of extracellular microparticles using ImageStream(X) imaging flow cytometry." Sci Rep 4: 5237.

Heinz, S., C. Benner, N. Spann, E. Bertolino, Y. C. Lin, P. Laslo, J. X. Cheng, C. Murre, H. Singh and C. K. Glass (2010). "Simple combinations of lineage-determining transcription factors prime cis-regulatory elements required for macrophage and B cell identities." Mol Cell 38(4): 576–589.

Hosokawa, H., J. Ungerback, X. Wang, M. Matsumoto, K. I. Nakayama, S. M. Cohen, T. Tanaka and E. V. Rothenberg (2018). "Transcription Factor PU.1 Represses and Activates Gene Expression in Early T Cells by Redirecting Partner Transcription Factor Binding." Immunity 48(6): 1119–1134 e1117.

Hunt, S. E., W. McLaren, L. Gil, A. Thormann, H. Schuilenburg, D. Sheppard, A. Parton, I. M. Armean, S. J. Trevanion, P. Flicek and F. Cunningham (2018). "Ensembl variation resources." Database (Oxford) 2018.

Jumper, J., R. Evans, A. Pritzel, T. Green, M. Figurnov, O. Ronneberger, K. Tunyasuvunakool, R. Bates, A. Zidek, A. Potapenko, A. Bridgland, C. Meyer, S. A. A. Kohl, A. J. Ballard, A. Cowie, B. Romera-Paredes, S. Nikolov, R. Jain, J. Adler, T. Back, S. Petersen, D. Reiman, E. Clancy, M. Zielinski, M. Steinegger, M. Pacholska, T. Berghammer, S. Bodenstein, D. Silver, O. Vinyals, A. W. Senior, K. Kavukcuoglu, P. Kohli and D. Hassabis (2021). "Highly accurate protein structure prediction with AlphaFold." Nature 596(7873): 583–589.

Karpurapu, M., X. Wang, J. Deng, H. Park, L. Xiao, R. T. Sadikot, R. S. Frey, U. A. Maus, G. Y. Park, E. W. Scott and J. W. Christman (2011). "Functional PU.1 in macrophages has a pivotal role in NF-kappaB activation and neutrophilic lung inflammation during endotoxemia." Blood 118(19): 5255–5266.

Klemsz, M. J. and R. A. Maki (1996). "Activation of transcription by PU. 1 requires both acidic and glutamine domains." Molecular and cellular biology 16(1): 390–397.

Korczmar, E. A., A. K. Bookstaver, E. Ober, A. N. Goldfarb, D. G. Tenen and B. Q. Trinh (2024). "Transcriptional Regulation of the Lineage-Determining Gene PU.1 in Normal and Malignant Hematopoiesis: Current Understanding and Therapeutic Perspective." Front Biosci (Schol Ed) 16(2): 10.

Korczmar, E. A., A. K. Bookstaver, E. Ober, A. N. Goldfarb, D. G. Tenen and B. Q. Trinh (2024). "Transcriptional Regulation of the Lineage-Determining Gene PU.1 in Normal and Malignant Hematopoiesis: Current Understanding and Therapeutic Perspective." Front. Biosci. 16(2).

Krueger, F. (2017). "Trim Galore: A wrapper script to automate quality and adapter trimming as well as quality control." 0.4.5, from https://www.bioinformatics.babraham.ac.uk/projects/trim_galore/.

Lam, M. T., H. Cho, H. P. Lesch, D. Gosselin, S. Heinz, Y. Tanaka-Oishi, C. Benner, M. U. Kaikkonen, A. S. Kim, M. Kosaka, C. Y. Lee, A. Watt, T. R. Grossman, M. G. Rosenfeld, R. M. Evans and C. K. Glass (2013). "Rev-Erbs repress macrophage gene expression by inhibiting enhancer-directed transcription." Nature 498(7455): 511–515.

Leddin, M., C. Perrod, M. Hoogenkamp, S. Ghani, S. Assi, S. Heinz, N. K. Wilson, G. Follows, J. Schonheit, L. Vockentanz, A. M. Mosammam, W. Chen, D. G. Tenen, D. R. Westhead, B. Gottgens, C. Bonifer and F. Rosenbauer (2011). "Two distinct auto-regulatory loops operate at the PU.1 locus in B cells and myeloid cells." Blood 117(10): 2827–2838.

Levantini, E., S. Lee, H. S. Radomska, C. J. Hetherington, M. Alberich-Jorda, G. Amabile, P. Zhang, D. A. Gonzalez, J. Zhang, D. S. Basseres, N. K. Wilson, S. Koschmieder, G. Huang, D. E. Zhang, A. K. Ebralidze, C. Bonifer, Y. Okuno, B. Gottgens and D. G. Tenen (2011). "RUNX1 regulates the CD34 gene in haematopoietic stem cells by mediating interactions with a distal regulatory element." EMBO J 30(19): 4059–4070.

Li, W., D. Notani, Q. Ma, B. Tanasa, E. Nunez, A. Y. Chen, D. Merkurjev, J. Zhang, K. Ohgi, X. Song, S. Oh, H. S. Kim, C. K. Glass and M. G. Rosenfeld (2013). "Functional roles of enhancer RNAs for oestrogen-dependent transcriptional activation." Nature 498(7455): 516–520.

Li, W., D. Notani and M. G. Rosenfeld (2016). "Enhancers as non-coding RNA transcription units: recent insights and future perspectives." Nat Rev Genet 17(4): 207–223.

Li, Y., Y. Okuno, P. Zhang, H. S. Radomska, H. Chen, H. Iwasaki, K. Akashi, M. J. Klemsz, S. R. McKercher, R. A. Maki and D. G. Tenen (2001). "Regulation of the PU.1 gene by distal elements." Blood 98(10): 2958–2965.

Liberzon, A., C. Birger, H. Thorvaldsdottir, M. Ghandi, J. P. Mesirov and P. Tamayo (2015). "The Molecular Signatures Database (MSigDB) hallmark gene set collection." Cell Syst 1(6): 417–425.

Lu, D., R. Nakagawa, S. Lazzaro, P. Staudacher, C. Abreu-Goodger, T. Henley, S. Boiani, R. Leyland, A. Galloway, S. Andrews, G. Butcher, S. L. Nutt, M. Turner and E. Vigorito (2014). "The miR-155-PU.1 axis acts on Pax5 to enable efficient terminal B cell differentiation." J Exp Med 211(11): 2183–2198.

Melo, C. A., J. Drost, P. J. Wijchers, H. van de Werken, E. de Wit, J. A. Oude Vrielink, R. Elkon, S. A. Melo, N. Leveille, R. Kalluri, W. de Laat and R. Agami (2013). "eRNAs are required for p53-dependent enhancer activity and gene transcription." Mol Cell 49(3): 524–535.

Mercer, T. R., M. E. Dinger and J. S. Mattick (2009). "Long non-coding RNAs: insights into functions." Nat Rev Genet 10(3): 155–159.

Mueller, B. U., T. Pabst, J. Fos, V. Petkovic, M. F. Fey, N. Asou, U. Buergi and D. G. Tenen (2006). "ATRA resolves the differentiation block in t(15;17) acute myeloid leukemia by restoring PU.1 expression." Blood 107(8): 3330–3338.

Natoli, G. and J. C. Andrau (2012). "Noncoding transcription at enhancers: general principles and functional models." Annu Rev Genet 46: 1–19.

Ng, C. E., T. Yokomizo, N. Yamashita, B. Cirovic, H. Jin, Z. Wen, Y. Ito and M. Osato (2010). "A Runx1 intronic enhancer marks hemogenic endothelial cells and hematopoietic stem cells." Stem Cells 28(10): 1869–1881.

Nurk, S., S. Koren, A. Rhie, M. Rautiainen, A. V. Bzikadze, A. Mikheenko, M. R. Vollger, N. Altemose, L. Uralsky, A. Gershman, S. Aganezov, S. J. Hoyt, M. Diekhans, G. A. Logsdon, M. Alonge, S. E. Antonarakis, M. Borchers, G. G. Bouffard, S. Y. Brooks, G. V. Caldas, N. C. Chen, H. Cheng, C. S. Chin, W. Chow, L. G. de Lima, P. C. Dishuck, R. Durbin, T. Dvorkina, I. T. Fiddes, G. Formenti, R. S. Fulton, A. Fungtammasan, E. Garrison, P. G. S. Grady, T. A. Graves-Lindsay, I. M. Hall, N. F. Hansen, G. A. Hartley, M. Haukness, K. Howe, M. W. Hunkapiller, C. Jain, M. Jain, E. D. Jarvis, P. Kerpedjiev, M. Kirsche, M. Kolmogorov, J. Korlach, M. Kremitzki, H. Li, V. V. Maduro, T. Marschall, A. M. McCartney, J. McDaniel, D. E. Miller, J. C. Mullikin, E. W. Myers, N. D. Olson, B. Paten, P. Peluso, P. A. Pevzner, D. Porubsky, T. Potapova, E. I. Rogaev, J. A. Rosenfeld, S. L. Salzberg, V. A. Schneider, F. J. Sedlazeck, K. Shafin, C. J. Shew, A. Shumate, Y. Sims, A. F. A. Smit, D. C. Soto, I. Sovic, J. M. Storer, A. Streets, B. A. Sullivan, F. Thibaud-Nissen, J. Torrance, J. Wagner, B. P. Walenz, A. Wenger, J. M. D. Wood, C. Xiao, S. M. Yan, A. C. Young, S. Zarate, U. Surti, R. C. McCoy, M. Y. Dennis, I. A. Alexandrov, J. L. Gerton, R. J. O’Neill, W. Timp, J. M. Zook, M. C. Schatz, E. E. Eichler, K. H. Miga and A. M. Phillippy (2022). "The complete sequence of a human genome." Science 376(6588): 44–53.

Patro, R., G. Duggal, M. I. Love, R. A. Irizarry and C. Kingsford (2017). "Salmon provides fast and bias-aware quantification of transcript expression." Nat Methods 14(4): 417–419.

Pekowska, A., T. Benoukraf, J. Zacarias-Cabeza, M. Belhocine, F. Koch, H. Holota, J. Imbert, J. C. Andrau, P. Ferrier and S. Spicuglia (2011). "H3K4 tri-methylation provides an epigenetic signature of active enhancers." EMBO J 30(20): 4198–4210.

Petrovick, M. S., S. W. Hiebert, A. D. Friedman, C. J. Hetherington, D. G. Tenen and D.-E. Zhang (1998). "Multiple functional domains of AML1: PU. 1 and C/EBPα synergize with different regions of AML1." Molecular and cellular biology 18(7): 3915–3925.

Piacenza, L., M. Trujillo and R. Radi (2019). "Reactive species and pathogen antioxidant networks during phagocytosis." J Exp Med 216(3): 501–516.

Ponting, C. P., P. L. Oliver and W. Reik (2009). "Evolution and functions of long noncoding RNAs." Cell 136(4): 629–641.

Qian, F., J. Deng, Y. G. Lee, J. Zhu, M. Karpurapu, S. Chung, J. N. Zheng, L. Xiao, G. Y. Park and J. W. Christman (2015). "The transcription factor PU.1 promotes alternative macrophage polarization and asthmatic airway inflammation." J Mol Cell Biol 7(6): 557–567.

Qiu, K., D. C. Vu, L. Wang, N. N. Nguyen, A. K. Bookstaver, K. Sol-Church, H. Li, T. N. Dinh, A. N. Goldfarb, D. G. Tenen and B. Q. Trinh (2024). "Chromatin structure and 3D architecture define the differential functions of PU.1 regulatory elements in blood cell lineages." Epigenetics Chromatin 17(1): 33.

Qiu, K., Vu, D., Wang, L., Bookstaver, A., Dinh, TN., Goldfarb, AN., Tenen, DG., Trinh, BQ (2024). Chromatin structure and 3D architecture define differential functions of PU.1 cis regulatory elements in human blood cell lineages, BioRxiv.

Ramirez, F., D. P. Ryan, B. Gruning, V. Bhardwaj, F. Kilpert, A. S. Richter, S. Heyne, F. Dundar and T. Manke (2016). "deepTools2: a next generation web server for deep-sequencing data analysis." Nucleic Acids Res 44(W1): W160–165.

Rosenbauer, F., K. Wagner, J. L. Kutok, H. Iwasaki, M. M. Le Beau, Y. Okuno, K. Akashi, S. Fiering and D. G. Tenen (2004). "Acute myeloid leukemia induced by graded reduction of a lineage-specific transcription factor, PU.1." Nat Genet 36(6): 624–630.

Simon, R., A. Lam, M. C. Li, M. Ngan, S. Menenzes and Y. Zhao (2007). "Analysis of gene expression data using BRB-ArrayTools." Cancer Inform 3: 11–17.

Staber, P. B., P. Zhang, M. Ye, R. S. Welner, C. Nombela-Arrieta, C. Bach, M. Kerenyi, B. A. Bartholdy, H. Zhang, M. Alberich-Jorda, S. Lee, H. Yang, F. Ng, J. Zhang, M. Leddin, L. E. Silberstein, G. Hoefler, S. H. Orkin, B. Gottgens, F. Rosenbauer, G. Huang and D. G. Tenen (2013). "Sustained PU.1 levels balance cell-cycle regulators to prevent exhaustion of adult hematopoietic stem cells." Mol Cell 49(5): 934–946.

Subramanian, A., P. Tamayo, V. K. Mootha, S. Mukherjee, B. L. Ebert, M. A. Gillette, A. Paulovich, S. L. Pomeroy, T. R. Golub, E. S. Lander and J. P. Mesirov (2005). "Gene set enrichment analysis: a knowledge-based approach for interpreting genome-wide expression profiles." Proc Natl Acad Sci U S A 102(43): 15545–15550.

Sundstrom, C. and K. Nilsson (1976). "Establishment and characterization of a human histiocytic lymphoma cell line (U-937)." Int J Cancer 17(5): 565–577.

Tenen, D. G. (2003). "Disruption of differentiation in human cancer: AML shows the way." Nat Rev Cancer 3(2): 89–101.

Trinh, B. Q., N. Barengo, S. B. Kim, J. S. Lee, P. A. Zweidler-McKay and H. Naora (2015). "The homeobox gene DLX4 regulates erythro-megakaryocytic differentiation by stimulating IL-1beta and NF-kappaB signaling." J Cell Sci 128(16): 3055–3067.

Trinh, B. Q., N. Barengo and H. Naora (2011). "Homeodomain protein DLX4 counteracts key transcriptional control mechanisms of the TGF-beta cytostatic program and blocks the antiproliferative effect of TGF-beta." Oncogene 30(24): 2718–2729.

Trinh, B. Q., S. Ummarino, Y. Zhang, A. K. Ebralidze, M. A. Bassal, T. M. Nguyen, G. Heller, R. Coffey, D. E. Tenen, E. van der Kouwe, E. Fabiani, C. Gurnari, C. S. Wu, V. E. Angarica, H. Yang, S. Chen, H. Zhang, A. R. Thurm, F. Marchi, E. Levantini, P. B. Staber, P. Zhang, M. T. Voso, P. P. Pandolfi, S. S. Kobayashi, L. Chai, A. Di Ruscio and D. G. Tenen (2021). "Myeloid lncRNA LOUP mediates opposing regulatory effects of RUNX1 and RUNX1-ETO in t(8;21) AML." Blood 138(15): 1331–1344.

Uribe-Querol, E. and C. Rosales (2020). "Phagocytosis: Our Current Understanding of a Universal Biological Process." Front Immunol 11: 1066.

Uszczynska-Ratajczak, B., J. Lagarde, A. Frankish, R. Guigo and R. Johnson (2018). "Towards a complete map of the human long non-coding RNA transcriptome." Nat Rev Genet 19(9): 535–548.

Visel, A., S. Minovitsky, I. Dubchak and L. A. Pennacchio (2007). "VISTA Enhancer Browser--a database of tissue-specific human enhancers." Nucleic Acids Res 35(Database issue): D88–92.

Yan, Y., D. Zhang, P. Zhou, B. Li and S. Y. Huang (2017). "HDOCK: a web server for protein-protein and protein-DNA/RNA docking based on a hybrid strategy." Nucleic Acids Res 45(W1): W365–W373.

Zhao, Y., H. Teng, F. Yao, S. Yap, Y. Sun and L. Ma (2020). "Challenges and Strategies in Ascribing Functions to Long Noncoding RNAs." Cancers (Basel) 12(6).

