## Supplementary material for "An integrated DNA interactome and transcriptome profiling reveals a PU.1/enhancer RNA-mediated Feed-forward Regulatory Loop Regulating monocyte/macrophage development and innate immune functions": None

### SUPPLEMENTAL FIGURES

**Figure S1. Proportion of PU.1-E associated genes and PU.1 DEGs in HL-60 cells, related to figure 1.**

(A-B) Pie charts showing the proportions of protein-coding and noncoding genes. A) PU.1-E associated genes identified by interrogating PU.1-Es with the human gene annotation GRCh38.85 from the GENCODE Project. B) PU.1 DEGs identified by comparing RNA-seq transcript profiles in HL-60 cells between PU.1 siRNA and siControl (derived from GSE87055).

**Figure S2. *LOUP* induces Mono/MΦ differentiation, related to figure 4**

A) RT-qPCR analysis of *CD14* expression after *ex vivo* myeloid differentiation as described in Figure 4A.
B) Representative bright field images were acquired with a Zeiss LSM 880 Confocal microscope, 40x/1.30 magnification and the PMT detector camera.  pro-monocytic U937 cells were culture in 10nM PMA for 72h hours to induce Mono/MΦ differentiation.

C-D) RT-qPCR analysis of *CD11B* and *CD14* expression upon PMA-induced Mono/MΦ differentiation using U937 cells.

E-F) RT-qPCR analysis of *CD11B* and *CD14* expression. *LOUP*-depleted (sgLOUP) and control (sgControl) CRISPR/Cas9 U937 cells were treated with 10nM PMA for 72h hours to induce Mono/MΦ differentiation.

**<u>Figure S3. *LOUP* induces Mono/MΦ differentiation, related to figure 5</u>**

A) Volcano plot for transcriptome comparative analyses of global gene expression profiles of *LOUP*-depleted (sgLOUP) vs control (sgControl) U937 cells. Each point denotes a gene. Genes with at least 1.5-fold changes are shown in blue. Down-regulated genes by *LOUP* are shown in left and up-regulated -regulated genes by *LOUP* are shown in right. X-axis: Log2 fold change, Y-axis: -Log10(p-value).

B) Representative bright field images were acquired with a Zeiss LSM880 confocal microscope, 40x /1.30 magnification and the PMT detector. pro-monocytic U937 cells were culture in 10nM PMA for 72h hours to induce Mono/MΦ differentiation.

### QUANTITATION AND STATISTICAL ANALYSIS

Statistical analysis and data quantification were conducted using GraphPad Prism 10.0 software. Results are presented as mean values with standard deviation (SD). To determine the statistical significance of differences between two experimental groups, we employed the two-tailed Student's t-test, unless otherwise noted in the corresponding figure captions. A p-value of 0.05 or less was considered statistically significant.
